# Can ecological interactions drive evolutionary outcomes? Evidence from insect host shifts between parasitic and non-parasitic plants

**DOI:** 10.1101/2024.04.03.587887

**Authors:** B. Zelvelder, G.J. Kergoat, L. Benoit, T. Tsuchida, J. Haran, R. Allio

**Author notes:** **Corresponding authors:** Rémi Allio & Benjamin Zelvelder, 755 avenue du campus Agropolis, 34988, Montferrier-sur-lez, France. (GJK;). (LB;); (JH;). (TT;).

## Abstract

Phytophagous insects have specialized on virtually every plant lineage. Parasitic plants, however, are uncommon hosts. Among insects, only a single lineage of weevils, the Smicronychini, has successfully radiated on both parasitic and non-parasitic plants in a large panel of distantly related Asterid families. This unusual pattern suggests that major host plant shifts have occurred over the course of their diversification. Through the analysis of a phylogenomic dataset, we reconstruct for the first time their evolutionary history and ancestral host plant associations. Our results show that independent host plant shifts occurred both from parasitic to non-parasitic hosts and between distinct parasitic lineages. These results suggest that host shift mechanisms can be driven by ecological opportunities provided by plant-plant interactions. This first evidence of extreme insect host plant shifts apparently mediated by parasitic plant-plant interactions emphasizes the core importance of ecological interactions as driving forces behind insect host plant shifts.

## Introduction

Plant-insect interactions are one of the main components shaping terrestrial ecosystems and have therefore been extensively studied throughout all major fields of Ecology and Evolution. Indeed, studying these interactions offer a promising opportunity for bridging micro- and macroevolutionary processes [1]. To apprehend the fundamental processes associated with the evolution of plant-insect interactions, considering the notion of host repertoire (*sensu* [2]) is particularly relevant. Echoing the Hutchinsonian concept of the ecological niche [3] (see also [4]), phytophagous insects have a *fundamental* repertoire of suitable hosts and a *realized* repertoire of hosts actually used by the insect in its environment [2]. The *realized* repertoire of hosts can either remain the same (i.e. niche conservatism [5]) or be subject to evolution via: specialization (i.e. fewer hosts [6-7]), diet expansion (i.e. hosts acquisition without the loss of ancestral hosts [8]) or host plant shifts (i.e. hosts acquisition with the loss of ancestral hosts [9]; see also [2]). With this point of view, host plant shifts are often seen as the most extreme scenarios as they can be seen as a shift to a distinct biotic environmental space [10-11]. Adaptation to new host plants requires the adaptation to new structures, plant defenses, phenologies, micro-climates and so on (e.g. [12-18]). Altogether, this multidimensional nature of host plant shifts is *de facto* much more challenging to study than other processes. As a result, underlying forces that enable such changes in the host repertoire of phytophagous insects remain poorly understood.

The evolution of host repertoire in phytophagous insects’ phylogenies follow these general patterns: *(i)* host plant shifts are ubiquitous events at large taxonomic scales, *(ii)* phytophagous insect phylogenies do not match host plant phylogenies in most cases, and *(iii)* phylogenetic niche conservatism of host associations is more common at shallower taxonomic scales (see the reviews of [2,19-21]). The description of these patterns led to the formulation of several macroevolutionary scenarios such as the classical ‘escape & radiate’ arms race by Ehrlich and Raven [22], the oscillation hypothesis [23], and the musical chairs hypothesis [24]. All these scenarios posit that host plant shifts are diversification drivers, a claim that has often been demonstrated on empirical datasets [24-28] (but see [21,29-31]). Host plant shifts have also been shown to be at the origin of genetic divergence between lineages, sometimes leading to reproductive isolation and even to ecological speciation (see the review of [32]). Overall, the signature of host plant shift and adaptations to new hosts have been extensively studied, from complex genetic changes [26,28,33] (see also the reviews of [34-35]) to transcriptomic signatures [11,36-37] and even morphological plasticity [38]. However, although the importance of this phenomenon in the evolution of arthropods has been widely acknowledged, how host plant shifts occur in the first place and what drives the directionality of these events has received very little attention.

Under the right conditions, it appears that phytophagous insects can easily make host plant shifts, at least towards plants in their fundamental repertoire. For example, through a process called ecological fitting (*sensu* [39]), numerous host plant shifts have been documented following geographical opportunities, such as new potential hosts introduction or parasites invasion in a new geographical range [40-43]. For those species, it seems that the only barrier to host shifts was the host’s geographical range. Other barriers associated with the plant as a niche, such as the plant’s phenology [44-45], physical structures (e.g. [46-47]), detectability [48], and physiology or chemistry (e.g. [12,49]), can also be overcome and ultimately lead to host plant shifts. These barriers therefore hold an important role in the respective diversification of plants and insects [22,26,50-53]. The phylogenetic proximity between related plants that, on average, share most of these characteristics, may in turn explain the ubiquitous ‘minor’ host plant shifts observed at shallow taxonomic scales [39,54-55]. Nevertheless, several experiments have succeeded in forcing host shifts towards plants supposedly outside of the insect’s host repertoire, showing that, under the right conditions (i.e. a consistent exposition to a new host), host preference is a relatively labile trait (e.g. [38,56]). Therefore, the key factors that explain ‘major’ insect host plant shifts - between distantly related plant species - may rely on rarer ecological opportunities that attenuate those barriers, allowing insects to overcome them. Interestingly, one such opportunity can be found *in natura* thanks to the interactions between parasitic plants and their hosted phytophagous insects.

Parasitic plants (*ca*. 4,500 species from 28 families) have a unique heterotrophic lifestyle associated with the development of a multicellular organ - the haustorium - which allows them to absorb water and nutrients but also proteins, nucleotides or pathogens [57]. Phytophagous insects that live on parasitic plants are directly impacted by the nature of the compounds exchanged with the parasitized plants [58-59]. As a result, phytophagous insects that feed on parasitic plants evolve in unique circumstances of physiological proximity between phylogenetically distant plants. To date, only one group of insects is known to have radiated on parasitic plants, that being the weevils of the tribe Smicronychini (Coleoptera: Curculionoidea; [60-67]).

Smicronychini weevils form a widespread tribe known from all continents except Antarctica that consist of *ca*. 200 species classified in six genera, the genus *Smicronyx* being by far the most diverse with 156 described species [63]. They are essentially monophagous or narrowly oligophagous on closely related plants and their endophagous larvae develop either in stems, roots, seeds or fruits, sometimes inducing galls in their host (see figure 1 and Table S1 for details, [63]). Although this tribe is known for its association with parasitic plants, it has a diverse repertoire of phylogenetically unrelated hosts plants, belonging to four Asterid families: some Smicronychini species specifically develop in holoparasitic plants (obligatory parasites with no or little photosynthetic activity [68]), like dodders (*Cuscuta* - Convolvulaceae) or broomrapes (*Orobanche* - Orobanchaceae), others develop in hemiparasitic plants (obligatory or facultative parasites that still have some degree of photosynthetic activity [68]; such as *Euphrasia* or *Striga* - Orobanchaceae), while the rest develop in non-parasitic plants belonging to the Asteraceae, Convolvulaceae and Gentianaceae families (figure 1; Table S1; [69]). At least one species from North America, *Smicronyx quadrifer*, is known to feed in both parasitic and non-parasitic plants, starting its larval development in a *Cuscuta* species and completing it in one of a few Asteraceae hosts parasitized by the *Cuscuta* [61]. Based on this striking ecological observation and because Smicronychini have a remarkably diverse host repertoire compared to other tribes of Curculioninae [70-72], we hypothesize that the close physiological proximity between distantly related plants provided by parasitic plant-plant interactions may give unique host plant shift opportunities to insects feeding on them.

**Figure 1:**
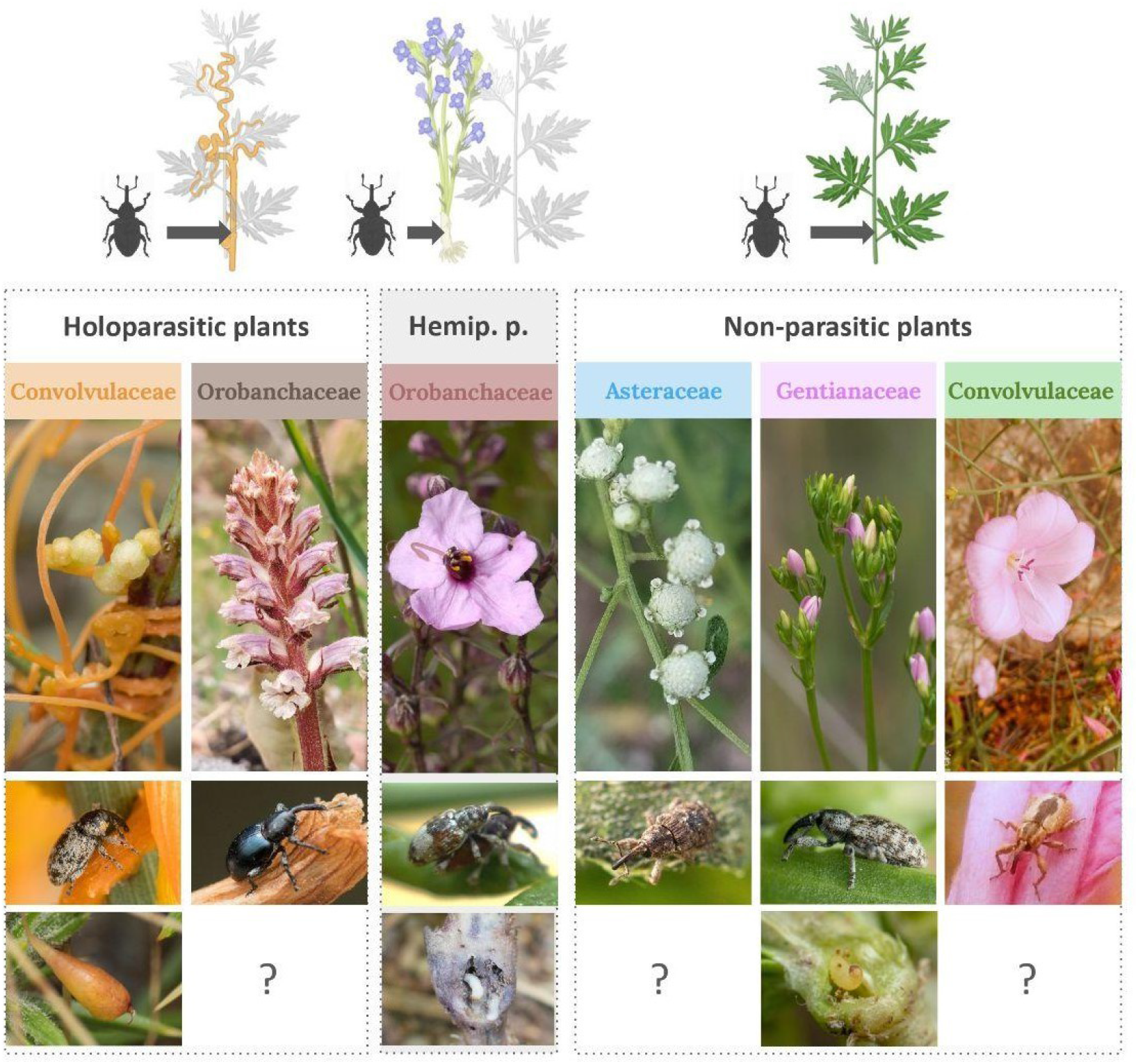
Host repertoire of Smicronychini weevils. From top to bottom: diagrams represent parasitic interactions between Smicronychini weevils and their host plants. Host plant families that make up the host repertoire of Smicronychini are shown below each corresponding diagram, with examples of associated weevils and known galls. Credits: B.Z., J.H., except for Orobanchaceae (Clémence Massard) and Asteraceae (@sinaloasilvestre, iNaturalist) pictures; diagrams with ©BioRender.

In this study, we address this hypothesis at the macroevolutionary scale by reconstructing ancestral host plant associations of the Smicronychini and propose a possible evolutionary scenario that may explain the host plant shift patterns inferred. We provide the first molecular phylogeny of the tribe following a two-stage approach. Firstly, we gathered and assembled hundreds of genome-wide markers using Anchored Hybrid Enrichment (AHE) capture, thus taking advantage of specimens held in museum collections to sample a representative dataset of the tribe. Secondly, we expanded our taxon sampling by grafting additional Smicronychini specimens/species for which mitochondrial markers (subunit I of the cytochrome *c* oxidase: COI) data was available. We then used the resulting phylogeny in combination with species-level ecological information to reconstruct the evolution of ancestral associations between Smicronychini and their host plants.

## Materials and methods

### Taxon sampling

The taxon sampling consists of 67 specimens intended for target capture (hereafter *AHE dataset*) and 205 specimens for which COI data were already available or newly generated (hereafter *COI dataset*).

*AHE dataset* - Smicronychini weevils were collected throughout ten years of field work carried out by J. Haran and by collecting specimens from Museum collections (Table S2). This sampling covers the Afrotropical, Palearctic and Oriental bioregions, representing all major genera encountered in this geographical range (i.e. *Afrosmicronyx, Sharpia, Smicronyx* and *Topelatus*), with the exception of the monospecific genus *Hedychrous*. Additional individuals were sampled in seven tribes (Anthonomini, Cionini, Curculionini, Derelomini, Erirhinini, Rhamphini and Styphlini) related to Smicronychini to serve as outgroups for phylogenetic inferences. These tribes were selected based on the most recent and comprehensive phylogenomic study of the Curculioninae subfamily [72].

*COI dataset* - COI sequences of individuals from the *AHE dataset* have been extracted using MitoFinder v1.4.1 [73]. When no COI sequence was found with MitoFinder, when available, COI sequences generated from the same sample or from different individuals of the same species were assigned to these individuals. This dataset allowed us to expand our taxon sampling and include taxa native to the Nearctic bioregion (poorly represented in the *AHE dataset*). Species from the genus *Promecotarsus* and the *Smicronyx* subgenera *Pachyphanes, Pseudosmicronyx* and *Smicronyx* were thus added, along with additional COI sequences of species from the *AHE dataset*. Those were either generated by our research group using specimens from museum collections or downloaded from GenBank (NCBI) or private and public data from BOLD Systems v4 [74] (see Table S2 for specimen details).

This sampling covers the entire morphological diversity and known host repertoires of the Smicronychini, considering both botanical families and the parasitic ecology of some of their hosts from the literature (Table S1), making it the most comprehensive sampling ever assembled for this tribe.

### Target capture and DNA sequencing

Specimens from the *AHE dataset* were prepared following the target enrichment protocol of [75]. Tissues from ethanol- and dry-preserved specimens were extracted non-destructively using the EZ-10 Spin Column Animal Genomic DNA Miniprep Kit (Biobasic Inc., Canada) with an overnight lysis step. DNA concentration was normalized (except for low quality samples) and mechanically fragmented to 300-600 bp fragments using a Bioruptor Pico (Diagenode; 15s ON, 90s OFF, eight cycles). Low quality DNA were then reconcentred with AMPure XP beads and libraries were constructed according to the NEBNext® Ultra II DNA Library Prep for Illumina® kit recommendations (New England Biolabs, Ipswich, MA, USA). Each library was individually labeled and subjected to 12 PCR cycles for final enrichment and pooled in equal quantities, grouping 24 libraries. The myBaits Hyb Capture for AHE kit (Arbor Biosciences, Ann Arbor, MI, USA) was applied to these two pools following the instructions of the manufacturer and assayed by qPCR. The resulting AHE library was sequenced paired-end, on a 2×150 bp SP line on an Illumina NovaSeq sequencer (Illumina, San Diego, CA, USA) at the GenomiX platform in Montpellier, France (MGX).

### Adapting probes to our dataset

The set of probes used during target enrichment was designed on 941 coding loci anchored in informative flanking regions within the genomes of distantly related coleopteran groups [76-77]. To improve the recovery of targeted loci on a shallower taxonomic level, we generated a new set of AHE reference sequences based on the complete genome of the boll weevil *Anthonomus grandis* (Curculionidae: Curculioninae; GCF_022605725.1, NCBI), the closest genome to Smicronychini at the time of this study (based on [72]). We mapped the entire probe set made available by Haddad et al. [76] on gene sequences of *A. grandis* using the *tblastx* command from BLAST+ 2.9.0 [78], which successfully retrieved all 941 genes targeted by the AHE probe set. Resulting matches were then filtered manually prior to phylogenomic analyses to remove 40 duplicated genes that mapped on multiple loci and 69 loci containing at least one insert more than 500 nucleotides long (out of 137 loci containing inserts in the putatively conserved region). The resulting set of 832 AHE reference sequences was further used to conduct assembly and alignment.

### AHE assemblies and alignments

A specific pipeline was developed to process AHE data (figure S1). Reads were first demultiplexed (i.e. reassigning reads to each individual on the basis of their tags used during target capture) with cutadapt v2.8 [79] and cleaned with Trimmomatic v0.39 [80]. Resulting reads were then assembled using a custom version of IBA [81]. This program follows an iterative approach to map reads on given reference sequences. Loci obtained from IBA assembly were then aligned to their corresponding AHE reference sequences using MAFFT v7.453 with *--add* and *--adjustdirectionaccurately* options [82-83]. We used the L-INS-I model, designed to be robust to divergent flanking regions around a core easier to align, which corresponds exactly to the AHE structure. To avoid any bias linked to missing data [84], we only retained loci represented by more than 70% of the individuals in our dataset, a value established as a good ratio between the quantity of loci retained and total percentage of missing data. Although AHE probes target coding regions, assembled sequences can be surrounded by non-coding flanking regions. Alignments were therefore cleaned up differently depending on their coding or non-coding nature. Firstly, partitions were defined based on the region targeted by the probes (custom script: *make_gapped_parts*.*py*). Secondly, using the annotation of the aforementioned genome of *A. grandis*, we were able to refine the boundaries of coding and non-coding regions more precisely (custom R script: *improve_parts*.*R*). This process led to the identification of potential overlapping sequences between AHE loci that were consequently pruned with cutadapt. Thirdly, coding and non-coding partitions were split using AMAS [85] and processed with the OMM_MACSE pipeline [86] or HMMCleaner [87] followed by PaulBlocks (provided by Paul Simion; to remove positions with more than 50% of missing data), respectively. All of these alignments were then concatenated into one supermatrix and translated into usable formats for phylogenetic inference using *seqCat*.*pl* and *seqConverter*.*pl*. This pipeline (scripts and details) is available at Zenodo (https://doi.org/10.5281/zenodo.15098685).

### Combined phylogenetic approach

#### Phylogenomic inferences based on AHE dataset

Phylogenetic inferences were conducted on the CIPRES Gateway cluster [88] within a Maximum Likelihood (ML) framework using IQTREE v2.3.2 [89-90]. The supermatrix was partitioned so that each flanking region, each inserted region and each coding position (figure 2B) was allowed to have its own evolution rate using the *-p* option [91]. Model estimation was systematically conducted using the greediest and most accurate method available in IQ-TREE using ModelFinder (*-MFP+MERGE* option) with *-rclusterf 20* to improve partition merging performance and the Bayesian information criterion (BIC) as a metric for model selection [92-93]. A perturbation level of heuristics of 0.2 (*-pers 0*.*2* option; recommended for a phylogenomic dataset according to IQTREE documentation) was implemented and symmetry tests were performed using the *--symtest-remove-bad* option [94]. The robustness of the nodes was estimated with ultra fast bootstraps (UFBS; [95]), with the *-bnni* option to compensate for model violations and the Shimodaira-Hasegawa approximate likelihood-ratio test (SH-aLRT; [96]), supposed to be more robust to the latter. A node was considered highly-supported for values of SH-aLRT ≥ 80 and UFBS ≥ 95% (following IQTREE authors’ recommendations).

**Figure 2:**
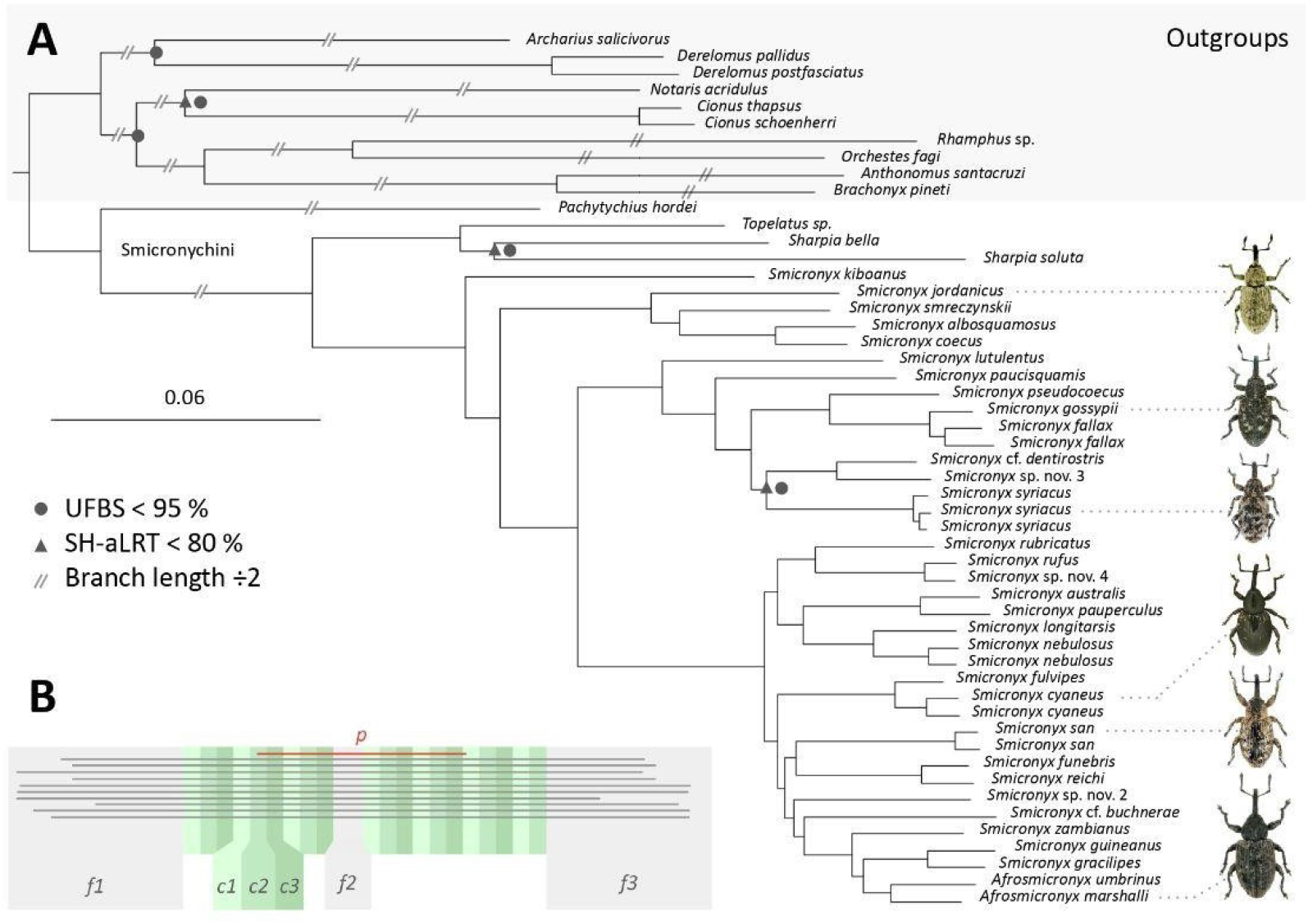
Phylogenomic tree based on the AHE dataset. **A**. Maximum likelihood tree of the AHE dataset. Nodes poorly supported (SH-aLRT < 80 or UFBS < 95%) are respectively highlighted with gray triangles and circles. For graphical purposes, some branches were shortened to half of their length (//). Pictures of weevils are linked to corresponding species. **B**. Partitioning scheme of a theoretical AHE loci. Flanking and inserted partitions (f1-3) are highlighted in grey and coding partitions (c1-3) are highlighted in green. The region targeted by the probes (P) is shown in red but does not appear in the final supermatrix. Credits for weevil pictures: J.H..

### Extending the phylogeny with the COI dataset

The tree obtained from the *AHE dataset* was pruned to contain only ingroup individuals (from the Smicronychini tribe). This tree was then used to constrain a second ML phylogenetic inference (*-g* option) on the *extended dataset* (i.e. including both *AHE* and *COI datasets*, excluding outgroups). This approach allows to densify the taxon sampling while retaining high confidence in the relationships at the species-level [97] (see [98-100] for empirical examples).

### Ancestral character state reconstructions

Ancestral character state reconstructions (ASR) of host plants were carried out using the R package *Phytools* v2.5.2 [101] to deal with polytomies and the R package *ape* v5.8 [102] to deal with unknown character states. Characters encoded in the main model correspond to known host plant families of Smicronychini (Asteraceae, Convolvulaceae, Gentianaceae and Orobanchaceae) associated with their parasitic interactions (non-parasitic, hemiparasitic and parasitic; figure 1; Table S1). Six categories and one unknown state were thus defined as follows: non-parasitic Asteraceae (A), non-parasitic Convolvulaceae (C), parasitic Convolvulaceae (pC), non-parasitic Gentianaceae (G), parasitic Orobanchaceae (pO) and hemiparasitic Orobanchaceae (hO). Alternative character states combinations were also tested using only host plant families or only parasitic types of host plants. ASR were carried out by comparing the likelihood of three Markov models (Mk), based on three different character state transition matrices, one with equal rates (ER), one with symmetrical rates (SYM) and one with all rates different (ARD). These models were then applied to the *extended dataset* phylogram to avoid biases linked to ultrametric trees [103-105], and reduced to one representative per species to limit ancestral trait estimation biases with the *fitMk* function. AIC values were compared using the *aic*.*w* function. Based on the best fitting model, each character state was assigned, at each internal node of the phylogeny, either a posterior probability using the *make*.*simmap* function over 1,000 simulations by a continuous-time reversible Markov process with *Phytools*, or a likelihood (L) directly estimated by the *ace* function with *ape*.

## Results

### Dataset filtration

*AHE dataset* - Prior to read assembly, the demultiplexing step highlighted the existence of sequences mapped to unused tag combinations due to their redundancy in target capture pools. These chimeric sequences originate from accidental “tag jumping” events described by [106], which occurred during the target capture protocol. Thus, three specimens with a number of reads falling below an arbitrary threshold of 2 Mpb were excluded from the dataset because their genetic information was considered indistinguishable from that of potential chimeras. For the same reason, two additional specimens were removed from the dataset as 0 ng/µL of DNA was detected following DNA extraction, even if DNA reads have been sequenced afterwards. Two cross-contamination events were also highlighted with MitoFinder and removed from the dataset. Finally, following the phylogenomic pipeline, eight individuals with less than 50% DNA information were filtered out, resulting in a final supermatrix of 52 taxa.

*COI dataset* - Out of 202 initial COI sequences, 153 were kept following filtration of sequences merged with missing COI from the AHE dataset, misidentifications, pseudogenes, contaminations and too short sequences.

### Phylogenomic pipeline

Following dataset filtration, the *extended dataset* consists of 195 specimens, representing 69 different species of Smicronychini. Interestingly, among the 62 individuals we know were sampled in dry museum collections, 40 were included in the final sampling (11 out of 19 in the *AHE dataset* and 29 out of 43 in the *COI dataset*). Specimens selected for the *AHE dataset* then followed our pipeline (figure S1). From 832 AHE reference sequences used for the iterative assembly 230 loci were filtered out mostly because of their low sampling coverage (less than 70% of the individual captured), resulting in 602 selected loci for the final supermatrix. The supermatrix built from the *AHE dataset* comprises 228,248 nucleotides from 52 individuals including outgroups. Interestingly, the elongation of the coding regions increases the coding part of the supermatrix from 50.3% to 61.3% of its total length. Overall, the concatenation of partitions carried out by ModelFinder reduced the number of partitions from 2703 to only 58, which is consistent with the high degree of coding partitions in the sequences obtained with AHE.

### Tree inferences

The phylogenomic tree of the *AHE dataset* is shown in figure 2A. It recovers the Smicronychini as a monophyletic tribe with maximum support, their most closely related outgroup is inferred to be from the genus *Pachytychius* (Curculioninae *uncert. sed*.; sometimes included in Smicronychini, e.g. by [107]). All but two nodes of the ingroup are inferred with maximum support (SH-aLRT > 99, UFBS > 98%). The genera *Sharpia* and *Topelatus* are recovered as sister to all remaining Smicronychini, but their monophyly is uncertain considering branch support (SH-aLRT = 5.3, UFBS = 53%). The most speciose genus *Smicronyx* is however found paraphyletic with the genus *Afrosmicronyx* in apical position embedded among *Smicronyx* species.

The phylogenomic tree of the *extended dataset*, consisting of both AHE and COI data, is shown in figure S2 and rerun pruned to one individual per species in figure 3. The addition of the genus *Promecotarsus* supports its monophyly (SH-aLRT = 90.1, UFBS = 95%) and reveals once again the paraphyly of *Smicronyx*, even though their relative position among *Smicronyx* species is poorly supported and variable between runs (SH-aLRT = 54.6, UFBS = 62%; figure S2-3). Consequently, eight major Smicronychini clades can be drawn from this topology: one non-parasitic Convolvulaceae-feeding *Sharpia* species, three independent clades of parasitic *Cuscuta*-feeding *Smicronyx* species, one non-parasitic

**Figure 3:**
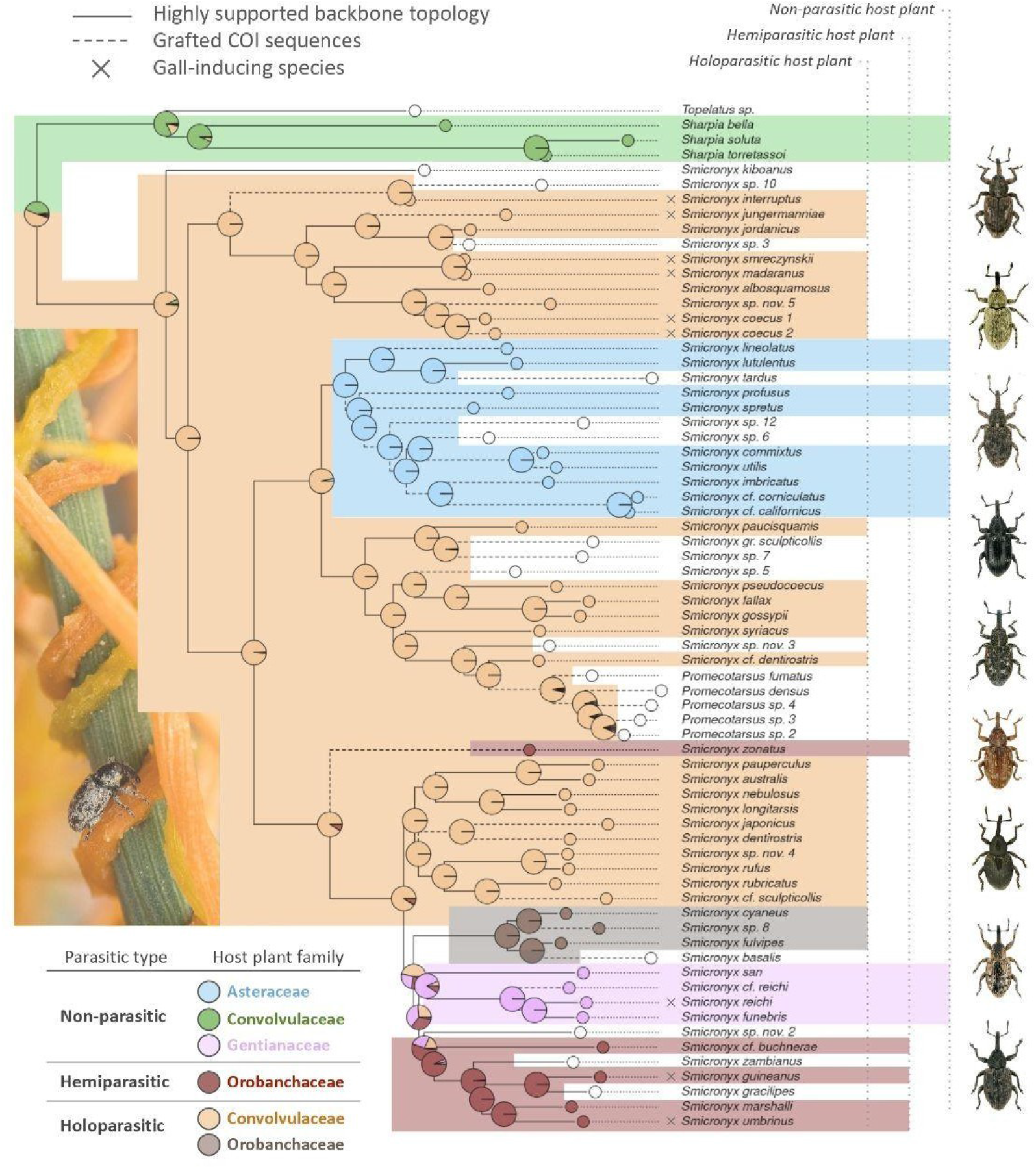
Ancestral host plant reconstruction (ASR) of Smicronychini weevils and their current ecologies. The phylogenetic tree topology corresponds to the extended dataset ML tree pruned to one branch per species. The AHE backbone tree is represented with plain branches while less robust COI data are represented with dashed branches. Reported gall-inducing species are marked with a “x” between branch tips and species names. Pie charts represented on each node of the phylogeny correspond to the estimated likelihood of each character state computed by the ace function. The colors of pie charts and background shading correspond to host plant character states used in the ASR as detailed in the bottom-left table. Colored shading represents the most probable ancestral state estimated at each node and left blank for unknown states and uncertain ancestral states. Photo credits: B.Z. & J.H.

Asteraceae-feeding *Smicronyx* species, two independent hemi- and parasitic Orobanchaceae-feeding clades of *Smicronyx* and *Afrosmicronyx* species and one clade of non-parasitic Gentianaceae-feeding *Smicronyx* species. However, surely because it mainly consists of COI grafted species, the Asteraceae feeding clade remains doubtful. Topology tests conducted between three topologies assuming (i) its paraphyly, (ii) its monophyly or (iii) its monophyly including *Promecotarsus* species revealed no significant differences in likelihood but ASR conducted over these alternative topologies differ slightly for this clade (see supplementary results on Zenodo at https://doi.org/10.5281/zenodo.15098685). Thus, the topology presented in figure 3 is the most parsimonious one of the equally likely topologies tested. In addition to *Promecotarsus* spp., four species remain difficult to assign to any of these clades, namely: *Topelatus* sp., *Smicronyx kiboanus, Smicronyx zonatus* and *Smicronyx* sp. nov. 2.

### Ancestral host plant estimations

ASR analyses of host-plant associations were conducted on the *extended dataset* pruned to one sample per species. Because chronograms are commonly used in ASR analyses, we additionally performed ASR on an ultrametric tree as a robustness check to assess whether our inferences were sensitive to tree type.ASR conducted on ultrametric trees or character state combinations restricted to plant families or parasitic types all agree with the main analysis presented in figure 3, that consists in a phylogram with plant families and parasitic types as character states (but see alternative ASR results on Zenodo at https://doi.org/10.5281/zenodo.15098685). The best fitting model is estimated to be the equal transition rate matrix (AICw: ER = 0.9998722; SYM = 0.0001278; ARD = 0). Results obtained from *Phytools* and *ape* packages being identical (Pearson’s correlation between matrices = 0.9999622), results of *ape* function that take into account unknown character states are presented in figure 3 to show additional species from the *extended dataset*. The most recent common ancestor of Smicronychini has an estimated likelihood of L_pC_ = 0.51 in favor of parasitic Convolvulaceae, L_C_ = 0.42 in favor of non-parasitic Convolvulaceae and less than 0.05 for every other character state. The parasitic Convolvulaceae character state is well supported as the ancestral state of all *Smicronyx* spp. *sensu lato* (including *Afrosmicronyx* and *Promecotarsus* regarding present analyses) with an estimated likelihood of L_pC_ = 0.92 or L_pC_ = 0.99 if we ignore *S. kiboanus*. Consequently, we infer several shifts from parasitic Convolvulaceae to other plant lineages such as non-parasitic Asteraceae (L_pC_ = 0.96) and parasitic Orobanchaceae (L_pC_ = 0.46, L_G_ = 0.25, L_hO_ = 0.23), without implying directional differences in transition rates or a causal driving role under the ER model. However, the ancestral host plant of weevils feeding on non-parasitic Gentianaceae and those feeding on hemiparasitic Orobanchaceae remains uncertain (L_pC_ = 0.26, L_G_ = 0.38, L_hO_ = 0.32).

## Discussion

This study provides the first robust phylogenetic hypothesis for the Smicronychini and reveals a remarkable pattern in the evolution of their host plant use. Phylogenomic inferences recover the monophyly of the tribe, but the paraphyly of the most speciose genus *Smicronyx* indicates that the current genus-level classification based on morphology is not satisfactory and requires a detailed re-investigation. That being said, the inferred *Smicronyx* clades consist of species with consistent morphologies and host plant associations, agreeing with the expectation of niche conservatism generally observed at a shallow taxonomic level in weevils (see the review of [70]).

The evolution of host plant preferences inferred herein reveals that the ancestral host plant of the tribe Smicronychini belongs with a high probability to the Convolvulaceae family. The most speciose lineage, represented by species belonging to the genus *Smicronyx* or closely related to it, are ancestrally associated with holoparasitic Convolvulaceae with great confidence. Interestingly, this lineage has experienced extreme secondary shifts toward phylogenetically distant botanical families encompassing both parasitic (Orobanchaceae) and non-parasitic Asterids (Asteraceae and perhaps Gentianaceae, both of which are known hosts of *Cuscuta* parasites). To our knowledge, this pattern is unique in phytophagous insects and is consistent with the hypothesis formulated in the introduction, which proposes that interactions between parasitic plants and their hosts may create ecological contexts conducive to major host plant shifts. The specific underlying molecular mechanism required to formally express this hypothesis has not been explored in the context of this study. However, the case of *S. quadrifer*, performing its development from a dodder to the host Asteraceae shows that such a transition is physically and physiologically possible. Based on this, we can formulate putative evolutionary explanations to the general host plant shift patterns that are observed. It has been shown that host-parasite interactions imply an intricate spatial proximity of tissues and allow exchanges of primary and secondary compounds between plants [108-109]. As such, the Smicronychini developing as larvae in the stems of parasitic plants are directly exposed to a physicochemical environment close to that of a dodder’s host. This shared environment can be seen as the ecological opportunity that led weevils feeding on dodders to evolve adaptations to their parasitic plant’s host, even though Asteraceae and Convolvulaceae have few secondary compounds in common (for Asteraceae see [110] ; for Convolvulaceae see [111]). Formulating hypotheses regarding shifts to other non parasitic plant lineages is more difficult, as it is unclear whether Gentianaceae feeding weevils shifted from parasitic Convolvulaceae or hemiparasitic Orobanchaceae.

Another intriguing host plant shift pattern in Smicronychini is the shift inferred between phylogenetically distant holoparasitic plant lineages, namely from Convolvulaceae to Orobanchaceae. This pattern suggests that a convergent parasitic lifestyle between phylogenetically distant lineages of plants could have promoted the shifts. One feature of parasitic plants that could explain this pattern is the existence of similar chemical compounds between dodders and broomrapes [112], like those involved in haustorium formation (e.g. [113-115]) that potentially facilitated the detection of these distant plants by one or a few Smicronychini lineages towards holoparasitic Orobanchaceae (although their detectability by Smicronychini have never been put to the test). Additionally, a second feature of parasitic plants is that they may share similar host plants and therefore show some degree of chemical similarity according to the proximity of their host plants. However, the host plant repertoire of Orobanchaceae-feeding weevils is not yet known with sufficient precision to corroborate this second hypothesis.

Phytophagous insect diversity has been widely associated with major shifts between distant plant lineages leading to bursts of diversification [19,22]. Yet, few papers have questioned how these major shifts occur in the first place. Although the patterns inferred in this study are specific to plant-plant parasitic interactions, the underlying scenario proposed here might provide valuable insights on the processes leading to major host plant shifts in phytophagous insects. From the insect perspective, plants with overlapping niches, for example with similar physiology, structure or chemistry, could provide ecological opportunities at the basis of host shifting events. In other words, overlapping plant niches can make up “facilitating bridges” that can be crossed by insects toward new potential hosts, as long as they are spatially and temporally consistent. Insects that shift from parasitic plants to their hosts make these opportunities more obvious in all of its multifaceted parts. It is likely that a large share of historical shifts has been driven by these kinds of opportunities, but the scarcity of ancestral data, intermediate states and our limited ability to link ecological niches in their multifaceted aspects to host plant shifts make it difficult to conceptualize. In this context, “minor” shifts that typically occur within a plant lineage - generally described as niche conservatism - may also derive from such ecological opportunities provided by the general relatedness (chemical, physiological, etc) between species in a given lineage. The decorrelation between phylogeny and niche relatedness inferred in the host repertoire of Smicronychini emphasizes the preeminence of ecological factors as prior components driving host shifts in phytophagous insects [54,116]. Similarly, many events of independent host plant shifts can be proposed as a result of ecological niche similarities at a broader scale. For example, the structural and physiological similarities in flower structures were suggested to be at the origin of major host shifts from monocots to dicots in derelomine flower weevils [117]. Another example is that of nematine sawflies (Hymenoptera: Tenthredinidae), where major host shifts occurred between phylogenetically distant, but ecologically similar hosts [118]. Altogether, such examples are consistent with a unified view of host shift mechanisms based on ecological opportunities arising from overlapping niches provided by plants.

## Ethics statement

The following services are acknowledged for providing permits: Western Cape Province (Western Cape Nature Conservation Board [permit No. CN44-30-4229], the Cape Research Centre [Republic of South African National Parks, CRC/2019-2020/012--2012/V1]), KwaZulu-Natal Province (Ezemvelo KZN Wildlife permits office, Collecting Permit KZN: OP1382-2019), the Direction de la faune de la chasse des Parcs et des Réserves (Niger: permit No. 1778177) and the Ministry of environment and sustainable development (Senegal: Authorization No. 231). Museum samples accession numbers can be found as part of supplementary material.

## Data accessibility statement

The dataset supporting this article is openly available in Zenodo at https://doi.org/10.5281/zenodo.15098685 [119]. Raw target capture data and assembled AHE results are publicly available in GenBank (NCBI) under the accession PRJNA1244829: https://www.ncbi.nlm.nih.gov/bioproject/?term=PRJNA1244829. COI sequences generated for the present study are publicly available on BOLD systems under the project name SCRYX and are accessible on Zenodo as well.

## Authors’ contributions

R.A., J.H., G.J.K. and B.Z. designed the study; J.H. collected data; L.B. and B.Z. conducted the experiments; B.Z. wrote the pipeline and performed all molecular analyses, supervised by R.A. and G.J.K.; B.Z. wrote the first draft; all authors substantially contributed to the writing of the final draft.

## Conflicts of interest

We declare we have no competing interests.

## AI declaration

We have not used AI-assisted technologies in creating this article.

## Fundings

This project was supported by the ANR Project ANR-21-CE32-0006 AgriBioDiv, coordinated by Christine Meynard.

## Acknowledgements

The following colleagues are acknowledged for providing some of the specimens used in this study: Robert Anderson (Canadian Museum of Nature, Gatineau, Canada), Patrice Bouchard (Ottawa Research and Development Centre, Ottawa, Canada), Lourdes Chamorro (USDA, Smithsonian Institution - National Museum of Natural History, Washington DC, USA), Desmond Edward Conlong (Stellenbosch University, South Africa), Madougou Garba (Direction Générale de la Protection des Végétaux, Ministère de l’Agriculture, Niamey, Niger), Hiroaki Kojima (Laboratory of Entomology, Tokyo University of Agriculture, Japan), Jean Pelletier (Monnaie, France) and Patrick Weil (Maison de la Nature et de l’Environnement, Pau, France). The analyses benefited from the GenoToul Bioinformatics platform (Toulouse, France) platform services. We also thank Christine Meynard for funding this project. This is contribution ISEM 2026-023 of the Institut des Sciences de l’Evolution de Montpellier.

## Notes

https://doi.org/10.5281/zenodo.15098685

### Competing Interest Statement

The authors have declared no competing interest.

### Summary of Updates

The overall tone of the manuscript was revised to avoid any misinterpretation of correlation as causation. Figure 3 was revised to improve clarity and to reflect minor adjustments in the methods used for ancestral character reconstruction. Supplemental files were updated.

