## Supplementary Figures for "Can ecological interactions drive evolutionary outcomes? Evidence from insect host shifts between parasitic and non-parasitic plants"

https://doi.org/10.1098/rspb.2026.0100

- Table S1 - Species represented in the extended dataset and their host plant associations
- Table S2 - Individual data and accession numbers
- Figure S1 - Flowchart of the phylogenomic pipeline developed to process AHE data
- Figure S2 - Extended dataset ML tree
- Figure S3 - Extended dataset ML tree pruned to one sample per species
- Link to Zenodo repository: https://zenodo.org/records/15098686

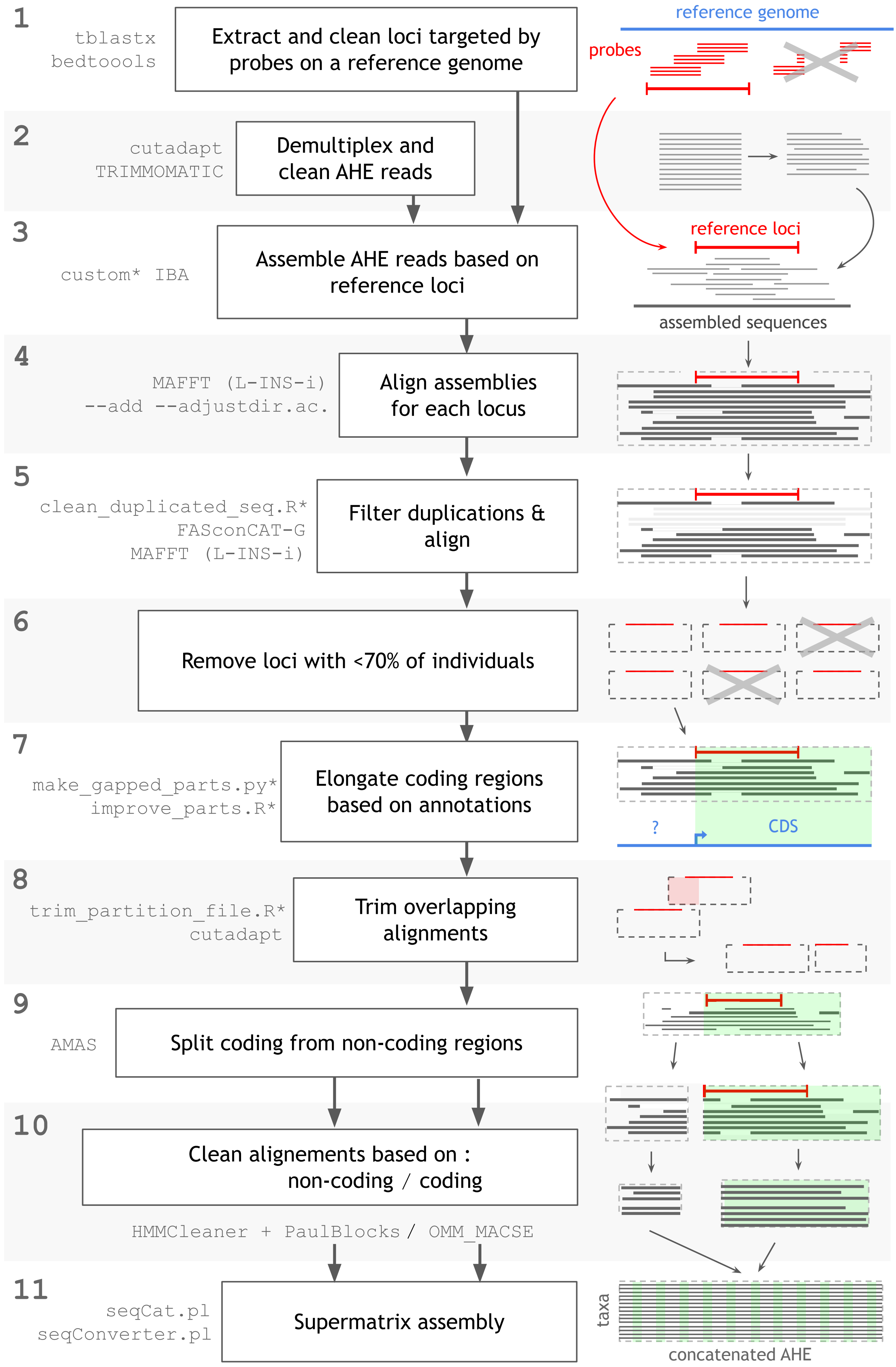


**Figure S1: Flowchart of the phylogenomic pipeline developed to process AHE data.** Each step, numbered from 1 to 10 as referenced in the main text, is represented, from raw reads to the complete supermatrix. External tools and custom scripts (marked with an *) are given, along with a diagram representing each step of the pipeline.

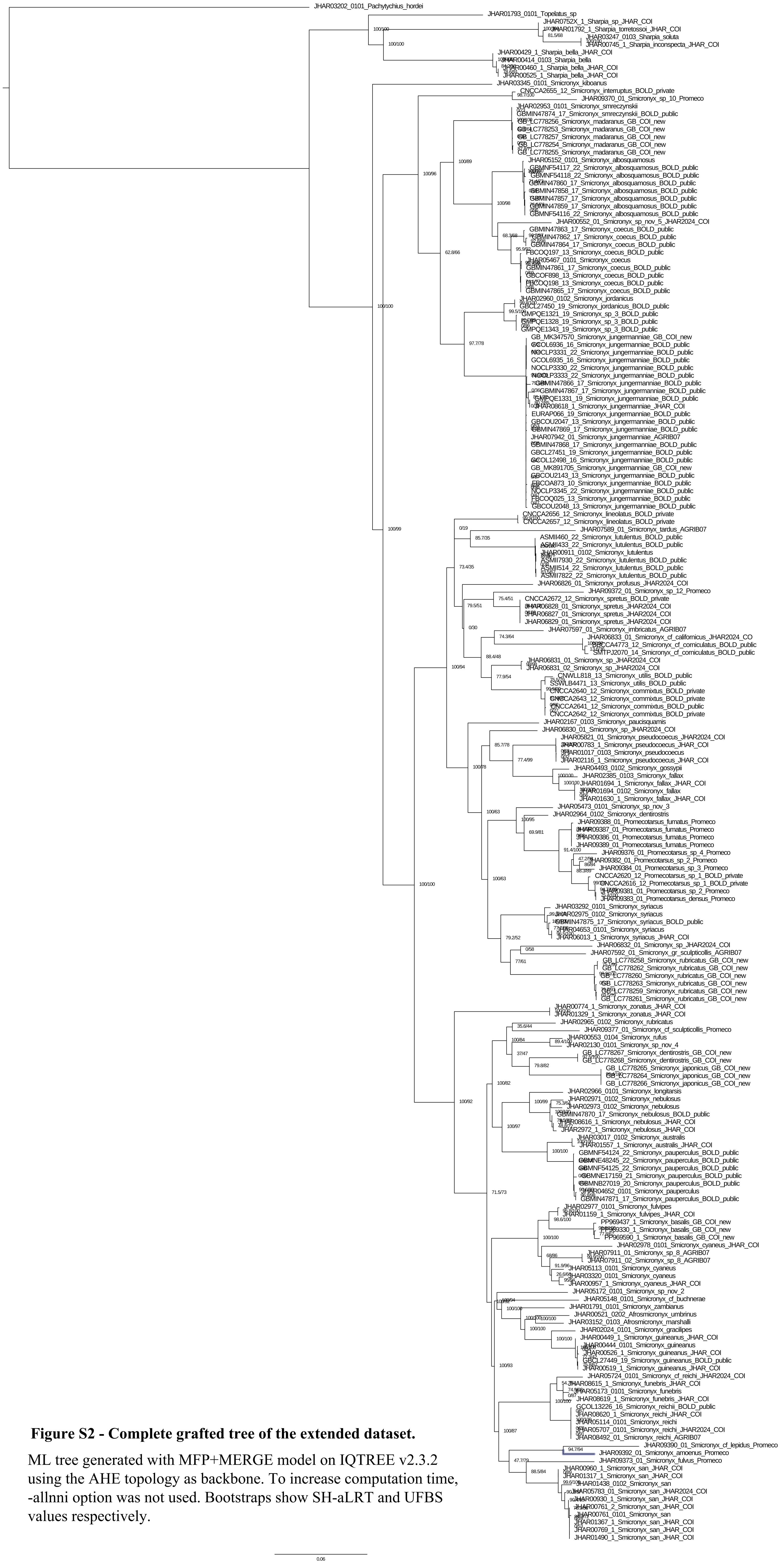

**Figure S2 - Complete grafted tree of the extended dataset.**

ML tree generated with MFP+MERGE model on IQTREE v2.3.2 using the AHE topology as backbone. To increase computation time, -allnni option was not used. Bootstraps show SH-aLRT and UFBS values respectively.

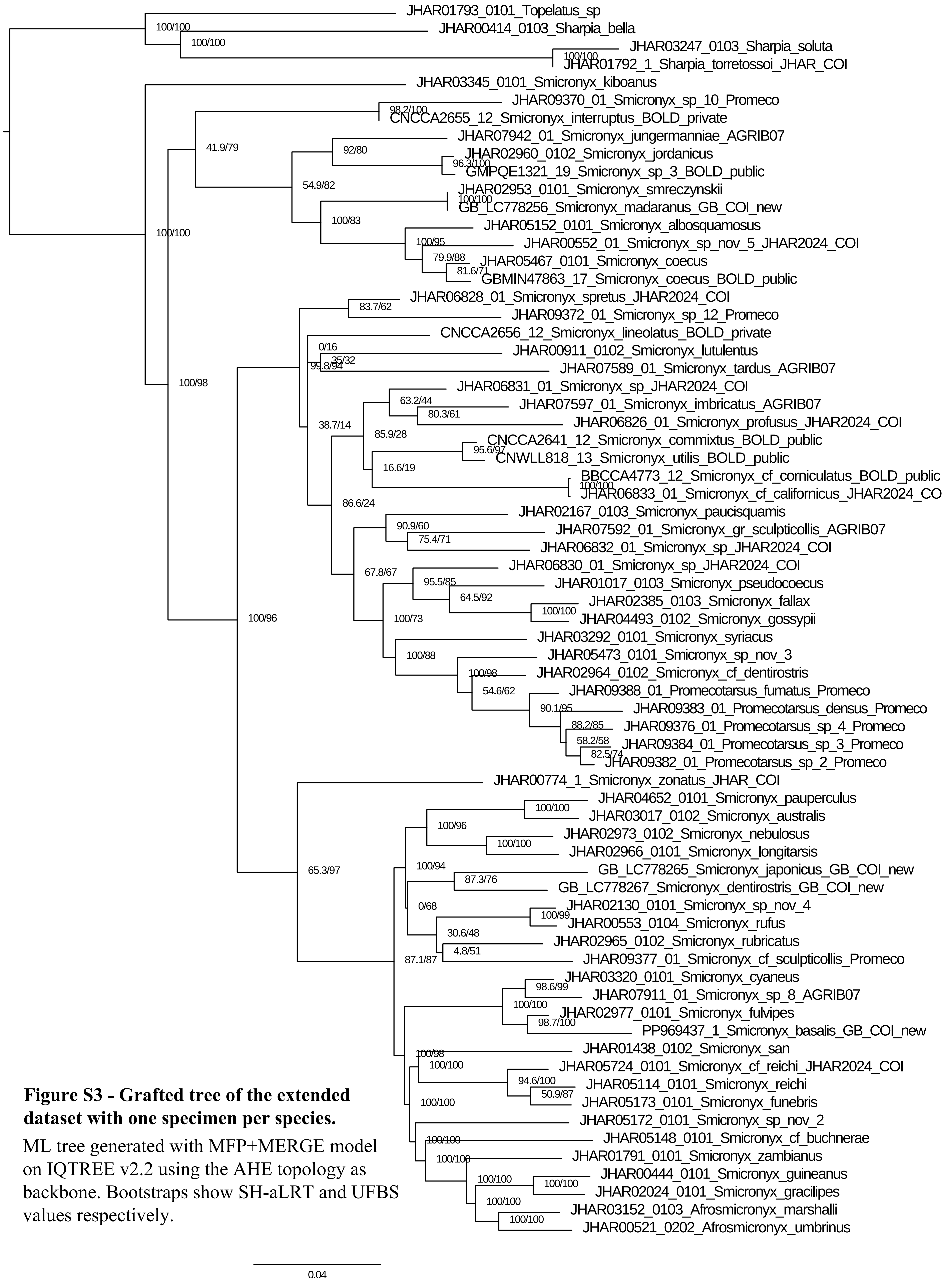


**Figure S3 - Grafted tree of the extended dataset with one specimen per species.**

ML tree generated with MFP+MERGE model on IQTREE v2.2 using the AHE topology as backbone. Bootstraps show SH-aLRT and UFBS values respectively.
